# Lysis buffer selection guidance for mass spectrometry-based global proteomics including studies on the intersection of signal transduction and metabolism

**DOI:** 10.1101/2024.02.19.580971

**Authors:** Barbara Helm, Pauline Hansen, Li Lai, Luisa Schwarzmüller, Simone M. Clas, Annika Richter, Max Ruwolt, Fan Liu, Dario Frey, Lorenza A. D’Alessandro, Wolf-Dieter Lehmann, Marcel Schilling, Dominic Helm, Dorothea Fiedler, Ursula Klingmüller

**Affiliations:** Division Systems Biology of Signal Transduction, German Cancer Research Center (DKFZ), Heidelberg, Germany; Translational Lung Research Centre Heidelberg, Member of the German Centre for Lung Research (DZL), Heidelberg, Germany; Proteomics Core Facility, German Cancer Research Center (DKFZ), Heidelberg, Germany; Leibniz-Forschungsinstitut für Molekulare Pharmakologie, Berlin, Germany; Institut für Chemie, Humboldt-Universität zu Berlin, Berlin, Germany

## Abstract

Prerequisite for a successful proteomics experiment is a high-quality lysis of the sample of interest, resulting in a large number of identified proteins as well as a high coverage of protein sequences. Therefore, the choice of suitable lysis conditions is crucial. Many buffers were previously employed in proteomics studies, yet a comprehensive comparison of lysate preparation conditions was so far missing. In this study, we compared the efficiency of four commonly used lysis buffers, containing the agents NP40, SDS, urea or GdnHCl, in four different types of biological samples (suspension and adherent cell lines, primary mouse cells and mouse liver tissue). After liquid chromatography-mass spectrometry (LC-MS) measurement and MaxQuant analysis, we compared chromatograms, intensities, number of identified proteins and the localization of the identified proteins. Overall, SDS emerged as the most reliable reagent, ensuring stable performance and reproducibility across diverse samples. Furthermore, our data advocated for a dual-sample lysis approach, including that the resulting pellet is lysed again after the initial lysis with a urea lysis buffer and subsequently both lysates are combined for a single LC-MS run to maximize the proteome coverage. However, none of the investigated lysis buffers proved to be superior in every category, indicating that the lysis buffer of choice depends on the proteins of interest and on the biological question. Further, we demonstrated with our systematic studies the establishment of conditions that allows to perform global proteomics and affinity purification-based interactome characterization from the same lysate. In sum our results provide guidance for the best-suited lysis buffer for mass spectrometry-based proteomics depending on the question of interest.

## Introduction

Intracellular signal transduction is orchestrated by proteins organized in networks and functional pathways, dynamically responding to environmental changes and cellular demands. Protein abundance has been shown to play a central role in shaping the complexity of signaling in these networks [1]. Although RNA transcript levels help to understand cellular processes by providing information on transcriptional changes, they fall short of providing a comprehensive understanding of the real-time state of entire pathways as the correlation of mRNA levels and protein abundance is usually low [2]. Therefore, achieving a systems-level perspective demands reproducible identification and quantification of proteins across the entire proteome.

Over the years, advances in mass spectrometry (MS)-based proteomics have transformed our ability to identify and, most importantly, quantify thousands of proteins in a single study [3, 4]. Cell lysis for protein extraction is an indispensable step in proteomic workflows, encompassing the extraction and denaturation of proteins from cells or tissues to enhance sensitivity in downstream detection, ensuring reliable quantification. Ideally, efficient and robust cell lysis leads to the isolation of all proteins not only from the cytoplasm but also from membranes and cell compartments. Thereby the complete amino acid sequence should stay intact. Unfolding the three-dimensional structure of proteins may facilitate access for downstream reagents and proteases in the sample preparation process.

However, the diverse nature of the properties and structure of proteins poses challenges to the lysis process. Particularly challenging is the lysis of transmembrane proteins, proteins in intracellular structures such as nuclei or proteins in complex with DNA, such as histones [5]. For instance, nuclear proteins, that are compartmentalized by the nuclei membrane and often bound to DNA, are challenging to lysate in substantial quantities. Another example is the hydrophobic nature of transmembrane proteins that can present an obstacle for complete proteins lysis. Therefore, the quality of quantitative data generated depends greatly on the characteristics and limitations of the lysis buffer of choice. While there have been previous reports on single-lysis solutions [6], developing a universally effective lysis buffer, capable of denaturing a diverse range of proteins from various subcellular compartments, remains challenging. Typically, lysis buffers rely on chaotropic denaturants such as urea or guanidine hydrochloride (GdnHCl), or on surfactants like the ionic detergent sodium dodecyl sulfate (SDS) or the non-ionic detergent Nonidet P-40 (NP40) [7, 8]. Bottom-up proteomics literature highlights the widespread use of urea over GdnHCl as the preferred chaotropic agent for classical in-solution digestion (ISD) [8, 9].

Although SDS has emerged as one of the most commonly used reagents for cell and tissue lysis, showcasing robust solubilization capabilities, including membrane proteins, detergents like SDS are not widely used in MS-based proteomics due to their potential interference with the downstream MS analyses. Even after cleanup, detergents may persist in the resulting peptide solutions, compromising the sensitivity and reliability of protein identification and quantification. Reports suggest that even low amounts of SDS (<0.01%) can potentially disrupt the reversed-phase separation of peptides and suppress ionization in electrospray techniques. The same holds true for urea-based lysis buffers, whose utilization necessitates additional steps in sample handling and cleanup strategies (i.e., StageTip), which employ C18 material in pipette tips or cartridges to desalt the recovered peptides [10]. These steps can introduce variability or peptide loss, potentially affecting the accuracy of quantitative measurements.

The implementation of the single-pot, solid-phase-enhanced sample-preparation (SP3) technology has played a major role in introducing alternative protocols for sample preparation and peptide clean-up prior to liquid chromatography-mass spectrometry LC-MS measurements [11]. SP3 takes advantage of a hydrophilic interaction mechanism between paramagnetic beads and proteins in a given sample, effectively eliminating any chaotropic reagent or detergents present in the lysis buffer. In this manner, SP3 is recognized as a more versatile and efficient approach to sample preparation, enhancing the adaptability of protocols and facilitating the use of reagents previously considered off-limits for MS-based proteomics. Despite previous studies successfully comparing widely used MS sample preparation methods (e.g., classical ISD and FASP) [12–14], there has been a notable gap in research, and a systematic evaluation of different lysis buffer reagents using a single sample preparation strategy has been missing.

Furthermore, lysates can be utilized to enrich for interaction partners before the analysis by MS. For these purposes it is essential to preserve the three-dimensional, native structure of proteins. Most widely employed reagents such as urea or SDS enhance solubility of proteins, while disrupting quaternary and tertiary protein structure, making their use unsuitable for the aforementioned applications [15]. However, NP40-containing lysis buffers, which are widely employed for immunoprecipitation and affinity binding studies, promise to not only be suitable for the analysis of transmembrane proteins by MS but also to facilitate affinity enrichment e.g. with a small molecule as bait, which enables high-throughput identification of protein binding partners by subsequent MS analysis [16–18]. Furkert *et al*. recently investigated highly negatively charged immobilized small molecule messengers, termed inositol polyphosphates, and observed reliable enrichment of inositol phosphate-binding proteins in two different mammalian cell lines [19], However, it remained unresolved whether these studies could be extended to cover not only intracellular but also interactions with integral membrane proteins.

Our analysis focuses on the quantitative differences introduced by different lysis buffers and highlights the need for tailored lysis buffer selection based on specific sample types providing valuable guidance for refining proteomic workflows.

## Experimental Procedures

### Cell culture

Human erythroid leukemia cell line AS-E2 cells were cultured at 37 °C, 5 % CO_2_ and 95 % rH in Iscove’s Modified Dulbecco’s Medium (Gibco), supplemented with 20 % (v/v) fetal calf serum (Gibco) and 1% (v/v) Penicillin-Streptomycin (Gibco). 2 U/mL erythropoietin (Erypo Epoetin alfa FS, Janssen) were freshly added after every splitting event.

Human hepatocellular carcinoma HepG2 cells were cultured at 37 °C, 5 % CO_2_ and 95 % rH in Dulbecco’s Modified Eagle’s Medium (Gibco), supplemented with 20 % (v/v) fetal calf serum (Gibco) and 1 % (v/v) Penicillin-Streptomycin (Gibco). For the experiment, 2×10^6^ cells were seeded in 6 well dishes (TPP) and cultured for two days.

### Isolation of mouse liver tissue and primary mouse hepatocytes

All animals used in the experiments were housed at the DKFZ animal facility under standard husbandry conditions in accordance with animal guidelines and regulations determined by German Cancer Research Center (DKFZ, Heidelberg). The governmental review committee of animal care of the state Baden-Württemberg approved the animal experiments described herein.

Mouse liver tissue was isolated from 8 weeks old male C57BL/6N mice after CO_2_ anesthesia. After carefully opening the abdominal cavity, the liver was resected and cut into 3×3 mm^3^ pieces. Liver pieces were immediately frozen in liquid nitrogen and subsequently powderized with a tissue homogenizer (B. Braun Micro-Dismembrator U Ball Mill).

Primary mouse hepatocytes were isolated from 11 weeks old male C57BL/6N mice according to a standard procedure [20, 21]. Anesthesia was carried out by intraperitoneal injection of 16 mg/kg bodyweight xylazine hydrochloride (2 % (w/v), Bayer HealthCare), 112.5 mg/kg ketamine hydrochloride (100 mg/mL, zoetis) and 15 mg/kg acepromazine (cp-pharma). After opening the abdominal cavity, the vena cava was cannulated with a 24G venous catheter. Then the liver was perfused with EGTA buffer (0.6% (w/v) glucose, 105 mM NaCl, 2.4 mM KCl, 1.2 mM KH_2_PO_4_, 26 mM Hepes, 490 µM L-glutamine (Gibco), 512 µM EGTA, 15 % (v/v) amino acid solution (70 mg/L L-alanine, 140 mg/L L-aspartic acid, 400 mg/L L-asparagine, 270 mg/L L-citrulline, 140 mg/L L-cysteine hydrochloride monohydrate, 1 g/L L-histidine monohydrochloride monohydrate, 1 g/L L-glutamic acid, 1 g/L L-glycine, 400 mg/L L-isoleucine, 800 mg/L L-leucine, 1.3 g/L L-lysine monohydrochloride, 550 mg/L L-methionine, 650 mg/L L-ornithine monohydrochloride, 550 mg/L L-phenylalanine, 550 mg/L L-proline, 650 mg/L L-serine, 1.35 g/L L-threonine, 650 mg/L L-tryptophane, 550 mg/L L-tyrosine, and 800 mg/L L-valine; pH 7.6); pH 8.3) with a flow rate of 8 mL/min for 7 min at 42°C. The vena portae was incised to allow buffer outflow. Subsequently, the liver was perfused with collagenase buffer (0.6% (w/v) glucose,105 mM NaCl, 2.3 mM KCl, 1.2 mM KH_2_PO_4_, 25 mM Hepes, 490 μM L-glutamine (Gibco), 5.3 mM CaCl_2_, 12 %(v/v) amino acid solution, 444 μg/mL collagenase type 1-A; pH 8.3) for 10 min at 42 °C with a flow rate of 8mL/min. After perfusion, the liver was removed and transferred to suspension buffer (0.6% (w/v) glucose, 105 mM NaCl, 2.4 mM KCl, 1.2 mM KH_2_PO_4_, 26 mM Hepes, 1 mM CaCl2, 0.4mM MgSO4, 0.2 % (w/v) BSA, 490 μM L-glutamine (Gibco), 15 % (v/v) amino acid solution; pH 7.6). By disrupting the liver capsule, hepatocytes are released. The resulting cell suspension was filtered through a 100 µm cell strainer and centrifuged at 50g for 5min at 4°C. Cells were resuspended in 10 mL adhesion medium (phenol red-free Williams E medium (PAN Biotech) supplemented with 10% (v/v) FCS (Gibco),0.1μM dexamethasone, 0.1% (v/v) insulin, 2 mM L-glutamine,1% (v/v) penicillin/streptomycin(Gibco)) and 2 million cells were seeded in 6cm collagen I-coated dish (BD BioCoat). Cells were cultivated at 37 °C, 5 % CO_2_ and 95 % rH for 4 h to allow adhesion, then washed by 37°C pre-warmed PBS three times. Cells were cultivated in 2 mL serum-depleted medium overnight until cell harvesting.

### Cell lysis

A two-step lysing procedure was applied to cell lysis. Four different lysis buffers were used for the first-step lysis. These were NP40-lysis buffer (1 % NP40, 0.1 % AEBSF, 0.1 % AP, RIPA salt (0.05 M Tris pH7.4, 0.15 M NaCl, 1mM EDTA pH8.8, 1 mg/mL sodium deoxycholate, 0.5 mM Na_3_VO_4_, 2.5 mM NaF)), SDS-lysis buffer (4 % SDS, 0.1 % AEBSF, 0.1 % AP, RIPA salt), urea-lysis buffer (4 M Urea, 50 mM NH_4_HCO_3_, 0.1 % AEBSF, 0.1 % AP), and GdnHCl-lysis buffer (6 M Guanidine hydrochloride,100 mM Tris 8.5, 0.1 % AEBSF, 0.1 % AP ). For the second lysis step, the same second-step lysis buffer was used (8 M Urea, 2 % SDS, 50 mM NH_4_HCO_3_, 0.1 % AEBSF, 0.1 % AP).

Liver tissue powder was lysed in 100 µl of the first-step lysis buffer. Each lysis buffer had three biological replicates. Samples were shaking for 30 minutes at 37 °C, then sonicated at 80 % amplitude for 60 seconds with 0.1-seconds-on / 0.5-seconds-off intervals, followed by boiling for 1 hour at 95 °C, (the boiling step is discarded for samples with lysis buffer containing urea). The samples were sonicated again and centrifuged at room temperature. The clear extract was collected as a supernatant lysate. The remaining pellet was lysed in 100 µl of the second-step lysis buffer and sonicated for 60 s, then centrifuged at room temperature. The cleared extract was collected and referred as pellet lysate. Lysates derived from the supernatant and pellet were mixed in 1:1 (v/v) ratio to generate the Mix lysate.

Primary mouse hepatocytes were harvested in 200 µl of the respective first-step lysis buffer. For AS-E2 cells, 1.5×10^6^ cells were collected, washed three times with PBS and finally lysed in 100 µL of the respective lysis buffer. For HepG2 cells, each well with a sample was washed three times with PBS before applying 250 µL of the respective lysis buffer to the well, afterwards the cells were harvested with a cell scraper.

The following procedure was applied to all samples from PMH, AS-E2 and HepG2 cells: Samples were incubated while rotating for 20 min at 4 °C, then sonicated at 75 % amplitude for 30 s with 0.1-seconds-on / 0.5-seconds-off intervals, followed by centrifugation at 4 °C. If lysis buffer contained SDS, centrifugation was performed at room temperature. The clear extract was collected as a supernatant lysate. The remaining pellet was lysed in 100 µl of the second-step lysis buffer and sonicated for 30 s then centrifuged at room temperature. The cleared extract was collected and referred as pellet lysate. Lysates derived from the supernatant and pellet were mixed in 1:1 (v/v) ratio to generate the Mix lysate.

### Sample preparation for mass spectrometry analysis

Sample preparation and protein digestion was performed according to the SP3 protocol adapted from [11]. Protein concentration of the lysates was determined by application of the BCA Protein Assay Kit (Pierce, Thermo Scientific). 20 µg protein per sample was used for further sample preparation. Reduction of disulfide bonds and alkylation was achieved with 40 mM tris-(2-carboxyethyl)-phosphine and 160 mM chloroacetamide at 95 °C for 5 minutes (samples containing urea were reduced and alkylated at 37 °C for 1 hour). SP3 magnetic beads were prepared as follows: per 10 samples 20 µL of each Sera-Mag Speed Beads A and B (GE Healthcare) were mixed 1:1, the suspension was placed on a magnetic rack until the supernatant appeared clear (approx. 1 min). While the supernatant was discarded, the beads were taken off the rack and were resuspended in water. This step was repeated three times. The final volume was adjusted 20 µL, from which 2 µL of the bead mixture was added to each sample. Protein binding to beads was induced by adding ethanol to a final concentration of 50 % (v/v) and incubating for 15 minutes, while shaking with 800 rpm at room temperature. After incubating, the samples were placed on a magnetic rack and the supernatant was removed after 1 min, subsequently beads were washed three times with 80 % ethanol. Afterwards, 75 µl of digestion buffer containing 0.8 µg Trypsin Gold (enzyme to protein ratio 1:25, Promega) in 100 mM triethylammonium bicarbonate (TEAB, Sigma Aldrich) was added to beads. The samples were sonicated in a water bath for 30 seconds, then incubated at 37 °C and 800 rpm overnight (12-16 hours) for protein digestion. Finally, the samples were placed on the magnetic rack for 1 min, the supernatant containing the digested proteins was collected and dried by vacuum centrifugation at 45 °C. Samples were stored at -20 °C until use.

### LC-MS/MS analysis

The digested peptides were resolved in 15 µl loading buffer containing 2 % acetonitrile and 0.1 % formic acid, then sonicated in water bath for 5 minutes and incubated at room temperature for 5 minutes. Chromatographic peptide separation was carried out using the Thermo EASY-nLC system extended over 120 minutes with a multi-stage gradient of solvent A (0.1 % formic acid in water) and solvent B (0.1 % formic acid in 80% acetonitrile): A 108-minutes active gradient was applied, where the solvent B increased from 8 % to 63 % with a flow rate of 350 nL/min. Peptide mixtures were analyzed by orbitrap mass spectrometer Q Exactive Plus (Thermo Scientific) connected to an electrospray ion source. Mass spectrometry data was obtained by one full scan with top 20 MS/MS scans. In Q Exactive Plus, the full scan MS spectra (350 to 1,500 m/z) was acquired with a resolution of 70,000 at m/z 200, a maximum injection time of 30 ms, and an AGC target value of 3×10^6^ charges. MS/MS spectra (200 to 2000) were acquired with a resolution of 17,500 at m/z 200, a maximum injection time of 50 ms, and an AGC target value of 10^5^ charges.

### Mass spectrometry data processing

For the lysis buffer comparison, raw mass spectrometry data was processed by MaxQuant (Version 1.6.3.3), as described by Cox J [22, 23], Proteins and peptides (minimal length 7 amino acids) identification was searched against the Mouse Uniprot FASTA database (2019-06-26) or Human Uniprot FASTA database (2019-03-31), respectively, using a target-decoy approach with a reversed database. Quantification of peptides and proteins was performed by MaxQuant with default settings. The original MaxQuant search result files will be made available on the ProteomeXchange consortium via PRIDE (https://www.ebi.ac.uk/pride/) repository. The file called “MaxQuant1” (Sup; Pellet; Mix) was used for the analyses shown in Fig. 2 to 4, while “MaxQuant2” (Frac treating Sup and Pellet as fractions; Mix) was used for Fig. 5 and 6.

**Figure 1:**
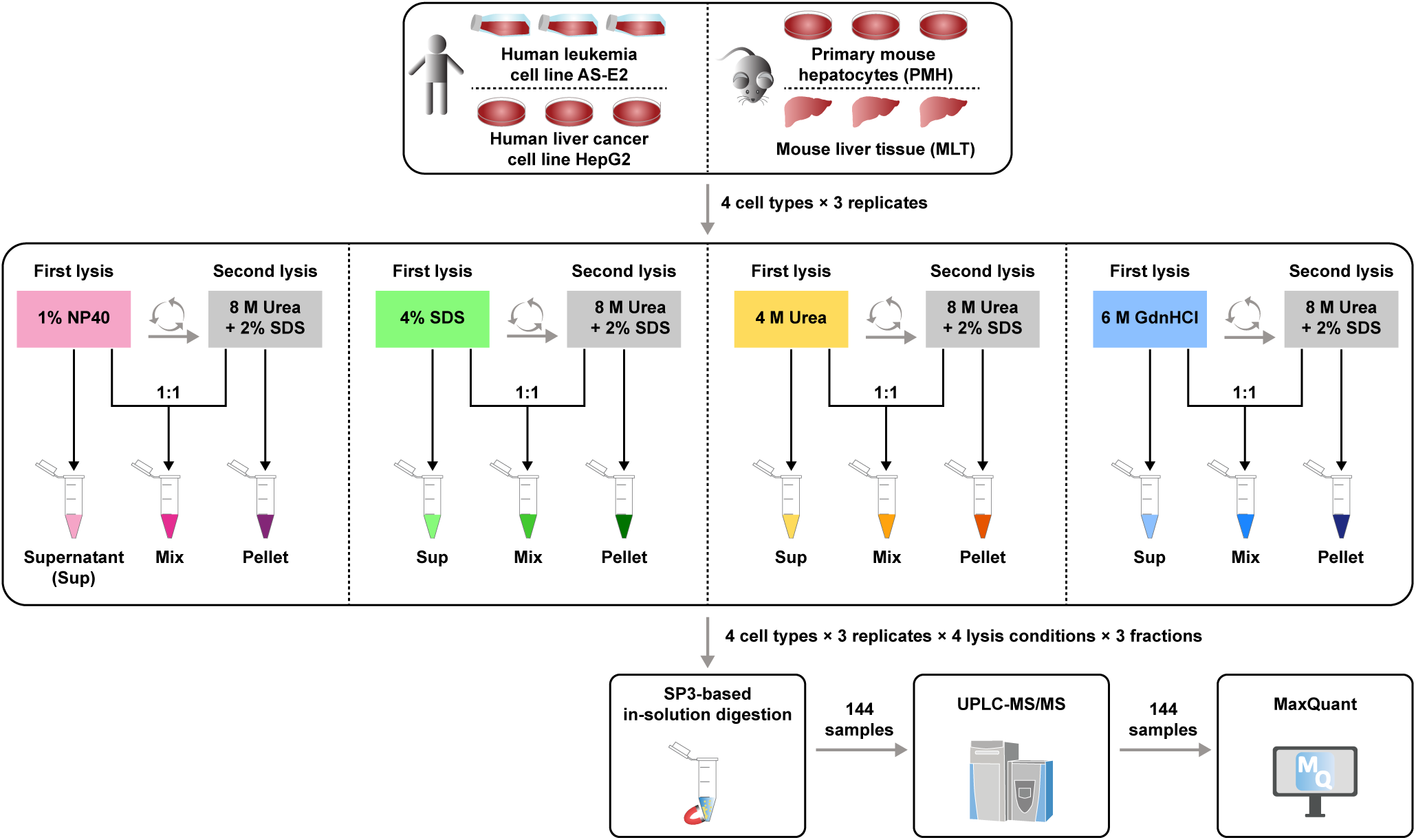
Experimental Workflow. A mass spectrometry-based quantitative proteomic strategy was employed for the systematic evaluation of various lysis buffer reagents during sample preparation. Solubilized proteomes from mouse liver tissue (MLT), mouse primary hepatocytes (PMH), adherent cells (HepG2), and suspension cells (AS-E2) were generated using different lysis buffers. The SP3 protocol was chosen as the primary method for sample preparation. The assessment included commonly used chaotropes and detergents at their typical concentrations: (i) 1% NP40, (ii) 4% SDS, (iii) 4 M urea, and (iv) 6M GnHCl. Each lysis step was followed by centrifugation, with the supernatant carefully retrieved. The remaining pellet was resolubilized in a lysis buffer containing 8 M urea and 2% SDS. This strategy allowed the separate or combined analysis (Mix) of distinct fractions (Supernatant or Pellet).

**Figure 2:**
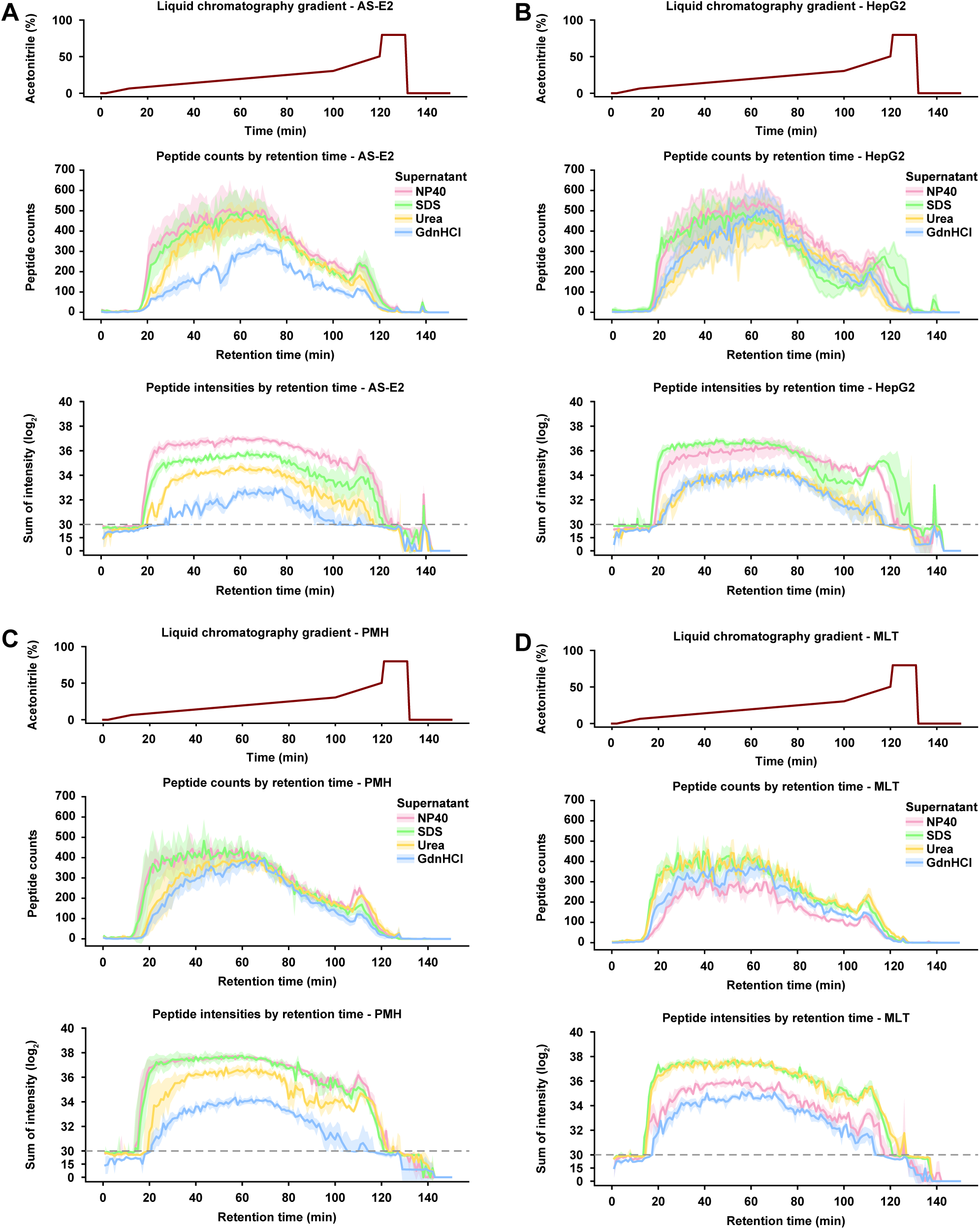
Liquid Chromatography and Peptide Elution Profiles. Nano-flow LC-MS/MS was performed by coupling an EASY-nLC 1000 system to an Orbitrap Q-Exactive Plus. A total of 20 µg of total protein was digested per sample, and the recovered peptides were dissolved in 15 µl 2% acetonitrile, 0.1% formic acid, and 5 µl were injected for each analysis. Peptides were directly delivered to an analytical column packed in-house with ReprosilGOLD C18, 1.9 µm resin, and separated using a 108 min active gradient from 8% to 63% of solvent B (0.1% formic acid, 80% acetonitrile; solvent A: 0.1% formic acid in water) at 350 nL/minute flow rate (top panels). The Orbitrap Q-Exactive Plus was operated in data-dependent mode (DDA), automatically switching between MS and MS2. The chromatograms from individual lysis buffers were overlaid for each supernatant, **(A)** AS-E2, **(B)** HepG2, **(C)** PMH, and **(D)** MLT, and visualized by retention time either as binned peptide counts or peptide intensities. Each line denotes the average of three replicates, with shadows representing the standard deviations.

**Figure 3:**
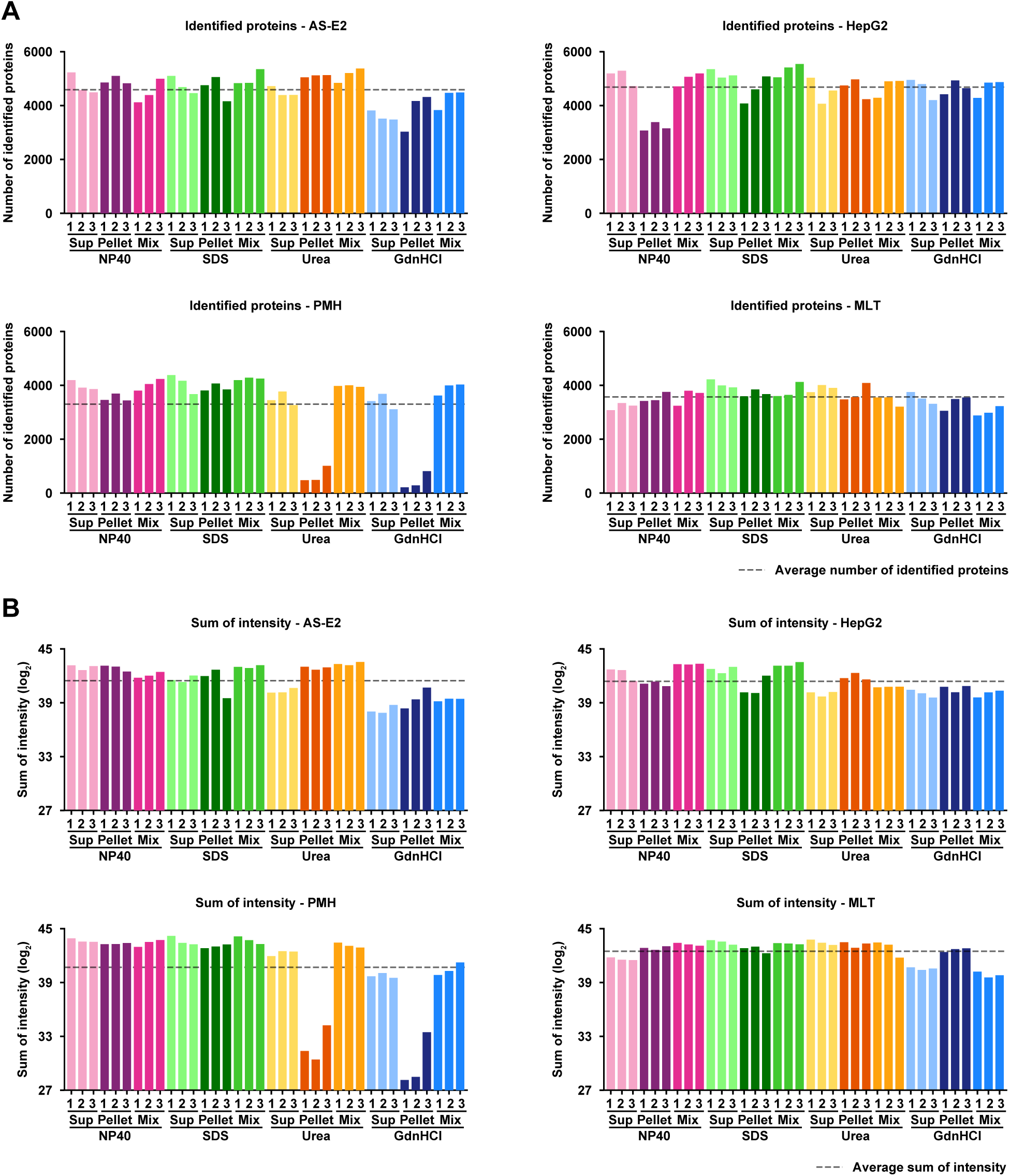
Reproducibility of Proteome Coverage. Samples of four sample types (AS-E2, HepG2, PMH, and MLT) were first lysed with RIPA buffer containing either 4M urea, 6M GnHCl, 4% SDS, or 1% NP40. After centrifugation the resulting supernatant was collected (Sup), while the pellet was treated with a lysis buffer containing 8M urea and 2% SDS in a second step. The extract was cleared by centrifugation (Pellet). Supernatant and Pellet samples as well as a 1:1 (v/v) mix of both (Mix) were analysed by LC-MS/MS as described before. Data analysis was performed with MaxQuant. **(A)** Bar graph of the number of identified proteins in every sample. Each replicate is depicted as one bar (n=3). The mean number of identified proteins over all samples is indicated by the dashed line. **(B)** Bar graph of the sum of all protein intensities. Each replicate is depicted as one bar (n=3). The mean sum of protein intensities over all samples is indicated by the dashed line.

**Figure 4:**
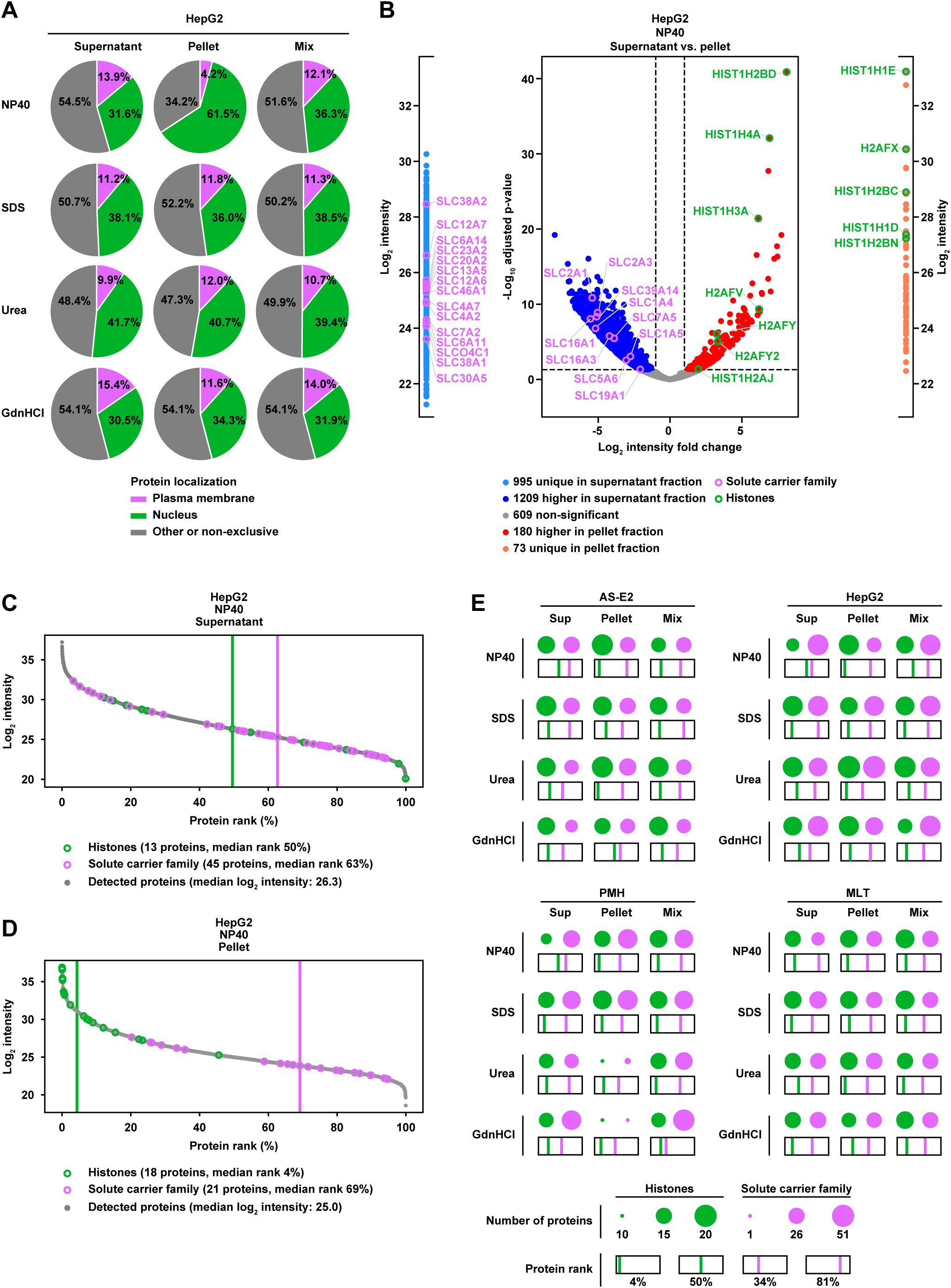
Distribution of Abundances of Nuclear and Membrane Proteins. According to their cellular localization all identified proteins were sorted in three categories: plasma membrane associated (pink), nuclear (green) and other/non-exclusive (grey). **(A)** Pie charts visualizing the percentage of protein intensities in the three categories, exemplified for HepG2. **(B)** Differential expression analysis depicted in a volcano plot comparing the abundance of proteins in Supernatant versus Pellet samples of HepG2 lysed with the NP40 lysis buffer. Average fold changes between Supernatant and Pellet and their statistical significance were calculated using the limma R package (cutoffs are set for a 2-fold change and an adjusted p-value < 0.05). Additionally, the intensities of unique proteins are shown at the sides of the volcano plot. Representative for plasma membrane-associated proteins, solute carrier family proteins (SLCs) are highlighted in pink, representative for nuclear proteins histones are highlighted in green. **(C)** and **(D):** Rank plots sorting all identified proteins in the Supernatant (C) or Pellet (D) of NP40-buffer lysed HepG2 cells according to their intensity from highest to lowest intensity. Vertical lines represent the median rank of the respective group. **(E)** Summary of SLC protein (pink) and histone (green) abundance in all lysis buffers, fractions and sample types. Dot size corresponds to the number of identified proteins of the respective group; the square underneath the dots indicates the mean protein rank of each group.

**Figure 5:**
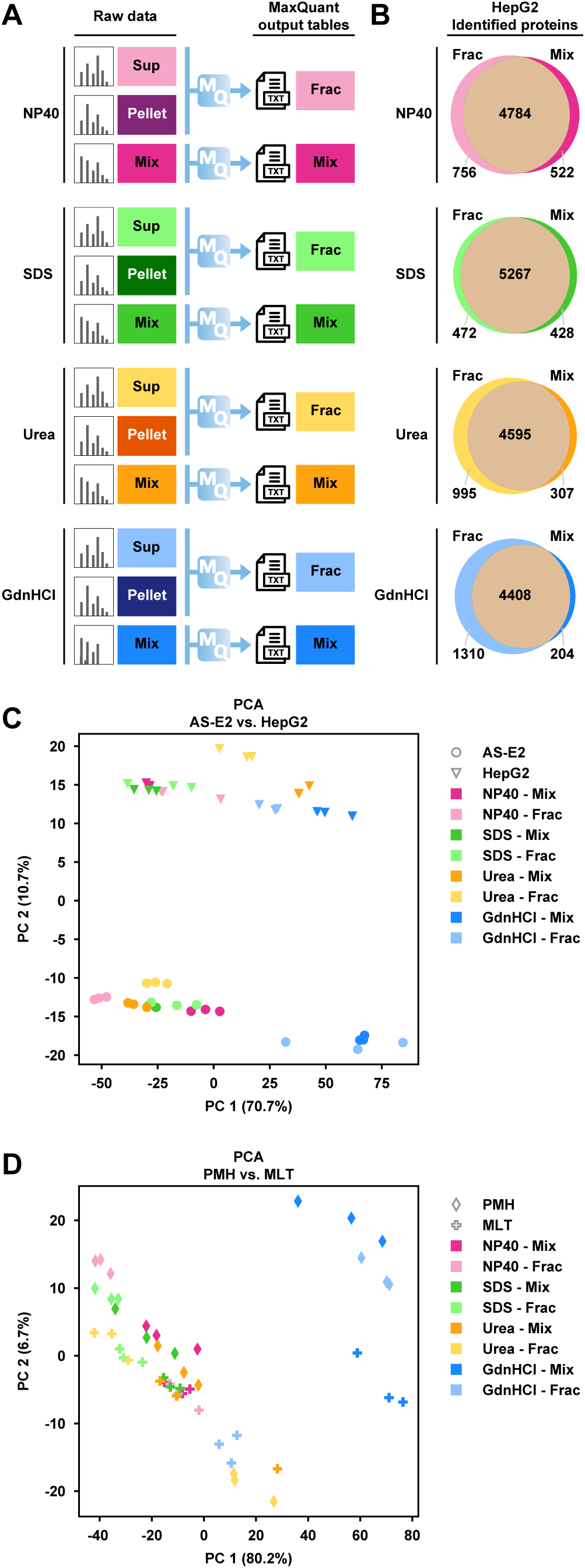
Consolidation of Supernatant and Pellet Fractionation. Comparing the Mix sample approach with combining the data of the Supernatant and Pellet *in silico*. **(A)** Workflow describing the generation of MaxQuant output tables used for further analysis. For each lysis buffer one output table was generated directly from the Mix sample results (Mix); a second output table was created combining the raw data from Supernatant and Pellet samples before processing them together in MaxQuant (Frac). **(B)** Venn diagrams depicting the overlap of identified proteins in Mix and Frac. Numbers represent the number of identified proteins in the group indicated. **(C)** and **(D)**: Principle component analysis plots to compare the maximum variance between AS-E2 and HepG2 samples (C), PMH and MLT (D).

**Figure 6:**
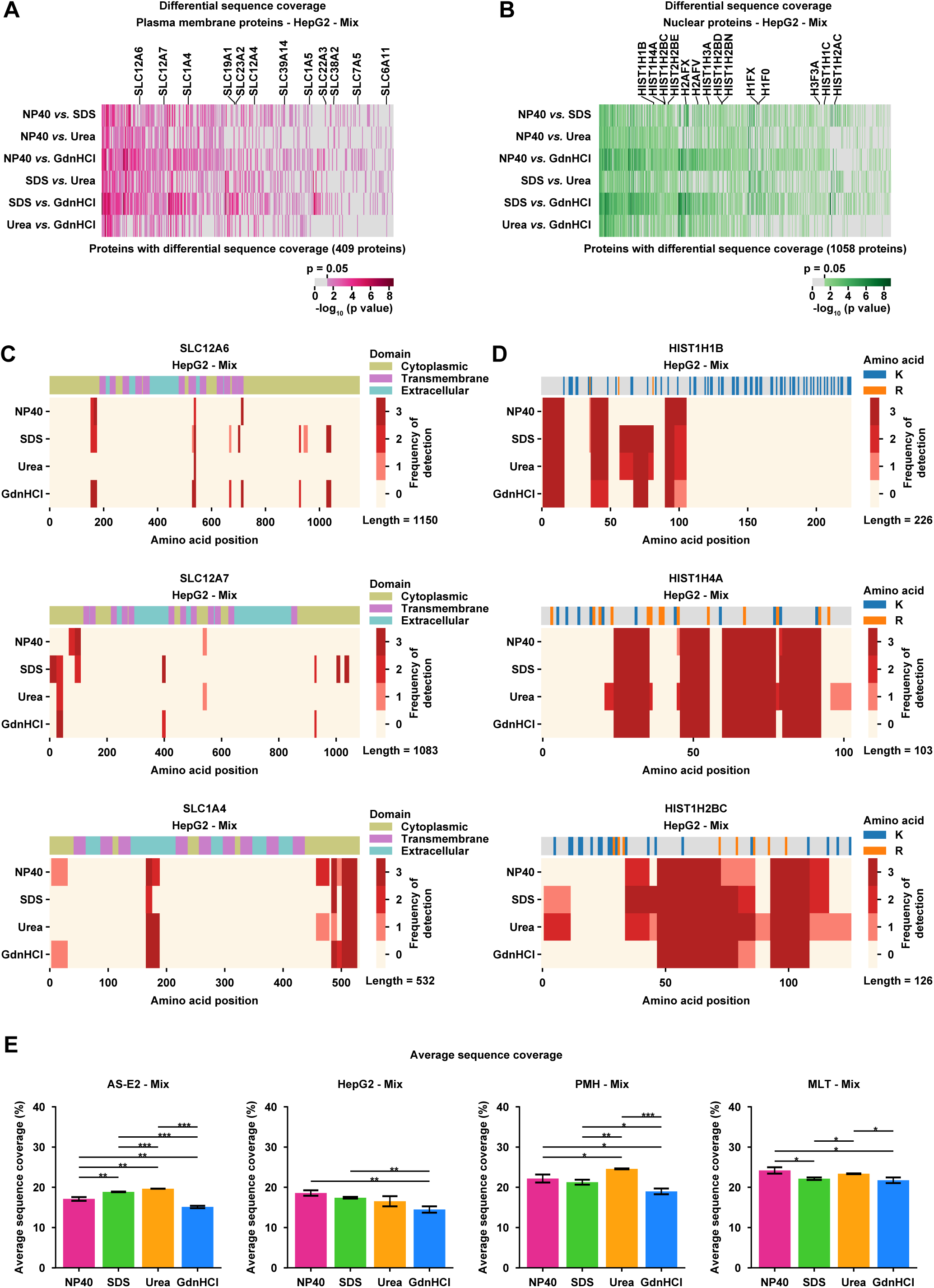
Differential sequence coverage and protein domain preference. **(A)** and **(B)**: Identification of proteins with statistically significant differential sequence coverage via limma testing between samples prepared with different lysis buffers. SLCs are highlighted among all proteins found to be statistically different (p adj. < 0.05) in sequence coverage among the plasma membrane proteins (A), while histones are highlighted for nuclear proteins (B). **(C)** Mapping of the detection frequency of peptides to the protein domains for the top three SLCs. Red scale indicating in how many replicates a respective peptide was detected. **(D)** Mapping of the detection frequency of peptides to the amino acid sequence of the protein for the top three SLCs. Red scale indicating in how many replicates a respective peptide was detected. **(E)** Average sequence coverage of all proteins detected in the mix samples of each sample type, comparing the coverage that could be achieved with the respective lysis buffer. Each bar represents the mean of three replicates, error bars indicate SD. *, p < 0.05, **, p < 0.01, ***, p < 0.001, t-test.

### Affinity enrichment for inositol polyphosphate interactome analysis

Multiscreen 96 well Plate, hydrophilic PVDF membrane was placed on the PlatePrep 96-well vacuum distributor and wells were washed with 70% EtOH (50 µL) followed by two washes with PBS (200 µL, 4° C). Streptavidin sepharose was added (120 µL) and washed with ice cold PBS (three times 200 µL). The bottom was dried with a clean wipe and the plate was positioned on a table top shaker at 4° C. 30 µL of a 1 mM biotin-1/3L-InsP_6_ solution was dissolved in 150 µL PBS and added to the wells. For control experiments 150 µL PBS was added and the plate was incubated at 300 rpm for 30 min. The plate was placed on the vacuum and washed with ice cold PBS (two times 200 µL) followed by one wash with RIPA lysis buffer (200 µL 25 mM Tris pH 7.6, 0.15 M NaCl, 1% sodium deoxycholate, 1% NP40 and 0.1% SDS.). The bottom of the plate was dried and it was placed on the table top shaker at 4° C. Cell lysate (150 µL, 1 mg/mL) was added and incubated at 300 rpm for 60 min. The unbound lysate was filtered off and the wells were quickly washed six times with RIPA lysis buffer at 4° C. The filter plate was placed on a conical 96 well-plate and competing InsP_6_ was added (50 µL, 5 mM in RIPA lysis buffer). The filter plate was incubated at 300 rpm for 15 min followed by centrifugation at 200 ×g for 2 min and the competition was repeated once more. The resulting affinity enriched lysates were used for proteomic sample preparation.

### SP3 tryptic digestion for inositol polyphosphate interactome analysis

Samples were reduced and alkylated at 37 °C for 1 hour adding chloroacetamide (40 mM final concentration) and tris(2-carboxyethyl)phosphine (5 mM final concentration). 2 µl of freshly prepared SP3 magnetic beads were mixed with samples [11]. Proteins binding to beads was obtained after adding 100 % acetonitrile to the final concentration of 50 % and incubating for 15 minutes at room temperature. The samples were subjected to a magnetic rack and the supernatant was removed, subsequently beads were washed three times with 70 % ethanol. 10 µl of digestion buffer containing 0.04 µg trypsin and 0.04 µg LysC in 50 mM TEAB buffer was added to beads and incubated at 37 °C overnight (12-16 hours) for protein digestion. The samples were subjected to the magnetic rack and washed twice with 200 µL acetonitrile. The beads were air dried and peptides were eluted adding 9 µL of 5% DMSO in Milli Q water and sonicating the solution for five minutes. The supernatant containing digested peptides was collected, dried under reduced pressure and stored at -20 °C.

### Liquid chromatography and mass spectrometry for inositol polyphosphate interactome analysis

After tryptic digest, peptide mixtures were separated by a reversed-phase chromatography (Dionex Ultimate 3000 NCS-3500RS Nano, Thermo Fisher Scientific) coupled on-line to an Orbitrap Exploris 480 mass spectrometer (Thermo Fisher Scientific, instrument control software version 4.2). The sample was first loaded on a PepMap C18 trap-column (75 µm ID x 50 mm length, 3 μm particle size, 100 Å pore size, Thermo Fisher Scientific) before reversed-phase separation on a 50 cm analytical column (in-house packed with Poroshell 120 EC-C18, 2.7 µm, Agilent Technologies) using an increasing acetonitrile gradient ranging from 4 – 80% over 120 min including column wash and equilibration at a flow rate of 250 nL/min. For standard protein identification and quantification, MS1 scans were acquired in a range of 375 to 1200 m/z with a resolution of 120,000 and an AGC target value of 3e^6^. Precursor ions with charge states 2-4 were isolated in a window of 1.6 m/z with a mass selecting quadrupole and 20 s dynamic exclusion time. MS2 scans were recorded with a resolution of 15,000, automatic scan range, AGC target and maximum injection time and an NCE of 30% in 2 s duty cycles.

### Identification, quantification and statistics of proteomics data for inositol polyphosphate interactome analysis

Raw data were searched using MaxQuant. Protein identification was performed with default settings. Search parameters included two missed tryptic cleavage sites, fixed cysteine carbamidomethyl modification, and variable methionine oxidation and N-terminal protein acetylation. Label-free quantification and match between runs were enabled. Database search was performed against the Human UniProt/Swiss-Prot database with common contaminants. The false discovery rate (FDR) was set to 1% at both peptide and protein level. The analysis of the quantitative proteomic data was carried out using Perseus. Proteins were filtered to exclude reverse database hits, potential contaminants, and proteins only identified by site. Data was imputed using Perseus default parameters.

## Results

### Establishing a Dual Cell Lysis Workflow for Unbiased Lysis Buffer Comparison

Cell lysis, encompassing the extraction and denaturation of proteins from cells or tissues, plays an essential role in proteomics analysis. While many protocols for the preparation of samples for mass spectrometric analysis have been described, systematic evaluation of the capacity of different lysis buffers in securing comprehensive representation of cellular proteins has been missing. Therefore, we carefully examined different lysis buffers, based on modifications of a lysis buffer widely employed in cell biology and mass spectrometry studies, to understand their impact on proteomics results. As depicted in the overview of our experimental workflow (**Fig.1)**, we selected three cellular models systems (i) the human leukemia cell line AS-E2, a cell line grown in suspension, (ii) the human liver cancer cell line HepG2, an adherent cell line, and (iii) primary mouse hepatocytes (PMH) and as model tissue mouse liver tissue (MLT). This diversity, spanning adherent, suspension, primary cells, and intact tissue comprising extracellular matrix was designed to uncover potential variations in lysis buffer performance across different cellular contexts, in order to systematically evaluate the relevance and applicability of our findings. Furthermore, by adopting SP3 as our primary sample preparation protocol, we utilized its versatility and efficiency to streamline the sample preparation process, offering a universally applicable approach that overcomes limitations present in conventional methods.

We strategically selected four distinct lysis buffers to represent varied biochemical properties. The buffers were based on the commonly used RIPA lysis buffer recipe as described in the Material & Methods part. To this buffer different lysing agents were added, representing those most commonly utilized in protein-based assays: 4 M urea, 6 M guanidine hydrochloride (GdnHCl, 4% sodium dodecyl sulfate (SDS), and 1% NP40. Their concentrations were chosen based on previously published studies [7, 24–26]. These lysis buffers were employed for the lysis of each sample type, which was called first lysis. The lysates were centrifuged, and the supernatant was collected (this lysate is called Supernatant sample in the following). Usually, the pellet of difficult to lyse components such as protein aggregates and extracellular matrix proteins, which is formed during centrifugation, is discarded. To test if we could improve coverage we resolubilized the pellets in a second lysis buffer containing 8M urea and 2% SDS. After centrifugation the resulting second supernatant was collected (this lysate is called Pellet sample in the following). Both Supernatant and Pellet samples were individually prepared and analyzed by mass spectrometry, allowing us to assess reproducibility and comparability of the results. In a subsequent step, the Supernatant and Pellet were recombined in a 1:1 (v/v) ratio for each sample, forming a mixed sample (in the following called Mix sample). Our approach aimed to reconstruct the protein composition of the original sample, leveraging the advantages of both the first and second lysis steps. To prepare the samples for the analysis by mass spectrometry, after lysis, the protein concentration was determined for each sample (Supernatant, Pellet, and Mix samples) using the bicinchoninic acid assay (BCA) assay. Protein concentration was normalized across samples, and 20 µg of total protein was used as input amount for digestion. The SP3 protocol was used for protein clean-up and digestion, and all samples underwent analysis in an EASY-nLC 1000 coupled to a QExactive Plus mass spectrometer operated in data-dependent acquisition (DDA) mode, and the acquired mass spectra were processed using MaxQuant. In total, four MaxQuant outputs were generated for the analysis of the three cellular model systems (HepG2, AS-E2 and PMH) and the tissue (MLT) and then used for unbiased statistical analyses. These studies revealed that our workflow is suitable to uncover distinct differences among the different lysis buffers for each sample type.

### Retention Time Profiles Demonstrate Reproducible, yet Lysis Buffer-specific Peptide Compositions

For an initial straightforward assessment of broad distinctions between the lysis buffers in the different cellular models and the model tissue, retention time profiles depict the peptide counts and the peptide intensities acquired by liquid chromatography after digestion. In our study, the peptide separation conducted on the EASY-nLC 1000 system extended over 120 minutes, with an active gradient from 12 to 120 min progressing from a 8 % to 63 % buffer composition containing 80% acetonitrile and 0.1% formic acid (**Fig. 2**, top panels in A, B, C and D). The chromatograms from individual lysis buffers were overlaid for each Supernatant sample, visualized either as binned peptide counts or peptide intensities by retention time. Each line denotes the average of three replicates, with shadows representing the standard deviations. Alignment of the averages indicates that observed effects originate from the lysis buffers, not alterations in the HPLC system. Moreover, equal amounts of peptides from each sample were used during analytical separation, ensuring that variations in intensity are attributed to lysis-buffer-related effects. Notably, the chromatograms resulting from urea, SDS, and NP40 lysis buffers are similar across all sample types, suggesting no general biases. No traces of SDS or NP40 persisted in the peptide solutions, as evidenced by the chromatogram consistency of the analysis of samples prepared by lysis buffers containing these detergents with the urea-containing lysis buffer profile for all sample types. GdnHCl presents a distinct profile, particularly in AS-E2 cells (**Fig. 2A**) and PMH (**Fig. 2C**), characterized by lower peptide counts and lower overall intensities compared to other lysis buffers. This effect of GdnHCl may impact its efficiency in specific cell types, underscoring the importance of tailoring lysis buffer selection based on cell characteristics.

### Proteome Coverage Demonstrates Cell-type Dependent Incomplete Protein Solubilization During the First Lysis

To identify the most effective lysis buffers for our diverse samples, we further evaluated the number of identified proteins (**Fig. 3A**). Using bar plots, we compared the number of identified proteins for each lysis buffer across all sample types. We observed a consistent count of over 4500 proteins in both suspension (AS-E2) and adherent cell lines (HepG2). The count decreased slightly to over 3500 for PMH and dropped to fewer than 3500 for mouse liver tissue. Noteworthy is the observed consistency among individual replicates for all lysis buffers, underscoring the robustness and reproducibility of protein identification. All lysis buffers performed similarly, yielding comparable numbers of total proteins identified in the supernatants. However, GdnHCl exhibited relatively lower performance, in particular for AS-E2 cells. Extending our analysis, we determined the total number of proteins identified in the pellets after the second lysis, employing 8M urea and 4% SDS. Surprisingly, a substantial number of proteins were also identified in the Pellet for almost all samples. This result highlights the importance of second lysis for quantitative studies and shows the efficiency of this procedure in rescuing proteins that would otherwise be discarded. However, for PMH following a first lysis with 4 M urea or 6 M GdnHCl, fewer proteins were recoverable from the pellet. We further extended our investigation to Mix samples, a combination of Supernatants and Pellets. This approach effectively equalized the numbers of identified proteins for all lysis buffers, mitigating potential variations in protein recovery. Given the substantial protein presence in the pellet, we asked whether the majority of intensity still originated from the Supernatant fraction, as lower intensities were expected in the Pellet fraction. To address this, we assessed the sum of protein intensities in each sample (**Fig. 3B**). A higher sum of intensities indicated superior protein extraction efficiency, showcasing the effectiveness of a specific lysis buffer. Notably, for MLT, HepG2 cells, and AS-E2 cells, the sum of intensities in the Pellet fraction of all lysis buffers equaled or exceeded those in the Supernatant, suggesting incomplete protein solubilization during the first lysis. Interestingly, for PMH, protein intensities in the Pellet fraction were lower than those in the Supernatant for urea and GdnHCl.

### Lysis Buffer-dependent Unequal Distributions of Nuclear and Membrane Proteins are Mitigated in the Combined Samples

Our results showed that a substantial number of proteins, both in terms of quantity and intensity, are found in the Pellet fractions of all tested lysis buffers in almost all sample types. Recognizing that discarding the pellet, a conventional practice, would entail a loss of numerous proteins and consequently lead to the loss of important qualitative and quantitative information. To gain a deeper understanding of potential losses in the pellet, we focused on investigating the classes of proteins that are overrepresented in the Supernatant or Pellet, aiming to disentangle potential disparities in protein composition and distribution between the two fractions. In particular, we evaluated two important classes of proteins that are challenging to be detect by mass spectrometry-based proteomics: transmembrane proteins and proteins derived from the nucleus. As exemplified by histones, nuclear proteins play a pivotal role in gene regulation, chromatin structure, and overall genomic stability. However, both the interaction with the DNA and the nuclear membrane have to be efficiently resolved to solubilize these proteins. On the other hand, transmembrane proteins, integral components of cellular membranes, are instrumental in mediating key processes such as signal transduction, transport, and cell-cell communication. The hydrophobic nature of their transmembrane domain renders this protein class particularly difficult to extract. Given their significance in cellular functions, neglecting either of these protein groups in further studies could result in an incomplete assessment of cellular processes determining dynamic behavior and cell fate decision. To conduct a comprehensive analysis, we categorized proteins in our dataset into three primary groups based on their cellular component annotation: (i) plasma membrane, (ii) nucleus, and (iii) other or non-exclusive. These categories were curated by manually extracting information from the Gene Ontology (GO), resulting in two main classes - membrane (comprising approximately 700 proteins) and nucleus (comprising around 3000 proteins). Following these categories, we employed a filtering process based on these protein categories. Subsequently, we calculated the sum of intensities for each category and normalized it by the total sum of intensities in the entire dataset. The resulting pie charts visualizes the relative distribution (percentage) of different protein categories for each lysis buffer, displaying the results for HepG2 cells as an example (**Fig. 4A**). Interestingly, the SDS and urea lysis buffers exhibited no discernible biases towards any specific protein class. The plasma membrane and nuclear protein distribution appeared consistent across the Supernatant, Pellet, and combined Mix. In contrast, the NP40 lysis buffer displayed a distinctive pattern, with a notable presence of plasma membrane proteins in the Supernatant fraction and an increased abundance of nuclear proteins in the pellet. Remarkably, the NP40 Mix sample closely resembled the distribution observed in

SDS and urea, suggesting that the original distribution is restored by combining Supernatant and Pellet. In the case of GdnHCl lysis, a distinctive trend emerged, indicating a slightly higher concentration of plasma membrane proteins and a reduced presence of nuclear proteins compared to other samples. Notably, the protein distribution in the GdnHCl mix revealed a persistent bias, suggesting that this particular lysis buffer exhibited a preference that cannot be fully rectified when combining Supernatant and Pellet fractions.

Next, to systematically evaluate the protein distribution across different fractions (Supernatant and Pellet), we examined two specific protein families, serving as distinctive markers for the plasma membrane (solute carrier -SLC-family) and nucleus (histones). The SLC proteins were chosen due to their intricate structure, characterized by multiple hydrophobic transmembrane alpha helices interconnected by hydrophilic intra- and extra-cellular loops. This complexity often poses challenges in achieving effective cell lysis and denaturation. On the other hand, histones, as nuclear proteins, are tightly bound to DNA through interactions between the backbone of their amino acids and the phosphodiester backbone of the DNA [27]. This close association may impede the representation of histones in proteomics analyses. Given the distinctive pattern observed in the NP40 lysis buffer, with a pronounced presence of plasma membrane proteins in the Supernatant fraction and an increased abundance of nuclear proteins in the pellet, we proceeded to perform a differential expression analysis, comparing the Supernatant and Pellet of this lysis buffer (**Fig. 4B**). Average fold changes between Supernatant and Pellet were calculated, and statistical significance was assessed using the limma R package, implemented into MSPypeline [28]. The significantly differentially expressed proteins are displayed by a volcano specifying an adjusted p-value (p<0.05) and a fold change (2-fold) as cutoffs. All proteins below these values were considered as non-significant (609 proteins). In total, 1209 proteins were more abundant in the Supernatant, and 180 proteins were more abundant in the Pellet, evidencing that in line with our expectation SLCs are more abundant in the Supernatant while histones are more abundant in the Pellet. Additionally, the intensities of the unique proteins of a group are shown at the side of the volcano plot and sorted according to their intensity. In total, 995 proteins were uniquely present in the Supernatant. Among these, 15 SLCs were observed. Conversely, 73 proteins were uniquely present in the Pellet, including 5 histones (**Fig. 4B**). To systematically investigate and compare the median intensities of SLCs and histones in both the Supernatant (**Fig. 4C**) and Pellet (**Fig. 4D**), we employed the MSPypeline to generate Rank plots. In this representation, the highest intensity in a given sample corresponds to rank 0 %, while the lowest intensity corresponds to rank 100 %. Consistent with our prior observations, SLCs emerged as a protein group with highest intensity in the Supernatant, while histones showcased high intensities in the Pellet. Summarizing these findings across all lysis buffers, fractions, and sample types encompassed in our study, we present the number of detected proteins in **Fig. 4E**. Pink dots signify SLCs, while green dots represent histones; the dot size corresponds to the number of SLCs or histones which were identified. The vertical bars indicate the protein rank, ranging from 1% on the left to 100% on the right, as in **Fig. 4C** and **Fig. 4D**. Our data reveals substantial differences between the Supernatant and Pellet across all lysis buffers. Interestingly, the mix demonstrated a remarkable capacity to restore not only the total number of proteins but also the abundances, as evident in the cases of SLCs and histones, except for GdnHCl. Notably, the SDS lysis buffer emerged as the most consistent lysis buffer across all fractions in our analysis (**Fig. 4E**).

### Consolidating Supernatant and Pellet Fractionation Ensures Enhanced Efficiency

Collectively, our findings strongly advocate for a dual-sample lysis approach, irrespective of the initial lysis buffer employed and the sample type. In the light of this, a pivotal question arises: Is it more advantageous to analyze the Supernatant and Pellet separately in two distinct LC-MS runs, later consolidating these fractions at the data level (e.g., prior to MaxQuant analysis) or should one combine the two fractions Supernatant and Pellet after the second lysis, opting for a single LC-MS run? This decision must consider the potential doubling of LC-MS time in the former method. In response, we carefully designed an experiment, as illustrated in **Fig. 5A**, where we analyzed either the Supernatant and Pellet fraction of each lysis buffer individually in the LC-MS, combining their data prior to MaxQuant analysis (Frac). The newly generated data of the two fractions (Frac data) was compared to the data of the Mix samples. **Fig. 5B** reveals a remarkable similarity in the number of identified proteins between both methods. Although distinct proteins are detected, the information collected from the analysis of the separate analysis of Supernatant and Pellet fraction does not appear to be substantially different from the analysis of the mix. To evaluate the potential introduction of bias during the sample mixing process, we conducted a principal component analysis (PCA), as shown in **Fig. 5C**. PC1 captures 70.7 % of variability, and separates samples mostly according to their lysis buffer type, specifically GdnHCl. In PC2, we observe a separation of the samples that clearly distinguishes samples based on their biological origin, AS-E2 and HepG2 cells. While the PCA recover the high reproducibility between both Mix and Frac samples, they also show that Frac and Mix conditions generally cluster together and contain similar information. Similar trends could be observed by comparing PMH and MLT (**Fig. 5D**). As expected, the hepatocytes more closely resemble the data of the liver tissue, compared to the two cell lines of different origins. In conclusion, our analysis suggested that measuring the Supernatant and Pellet fractions separately may be unnecessary. Instead, a more efficient approach could involve conducting two lysates and subsequently combining the two lysates. This strategy streamlines the process and maintains consistent analytical insights, as evidenced by the PCA results showcasing uniform behavior across various lysis buffers and fractions.

### Protein Sequence Coverage is Influenced by the Lysis Buffer Choice

While our attention has primarily been on protein identification numbers and intensities (**Fig. 2**), the sequence coverage of a protein holds equal significance. To ascertain whether the sequence coverage of individual proteins represented in the Mix samples of the different lysis buffers remains consistent across each lysis buffer or exhibits variations, we examined potential biases towards specific protein domains and display the results for HepG2 cells as examples. Considering the plasma membrane category and applying the limma test, we identified in pairwise comparisons of the data from the different lysis buffers proteins with statistically significant differences in sequence coverage between the four tested lysis buffers. The analysis revealed 409 proteins with differential sequence coverage, including 12 SLCs (**Fig. 6A**). The same assessment was performed for the nuclear category (**Fig. 6B**), identifying 1058 proteins, among which 15 were histones. We selected the top three proteins from each category to gain further insights and plotted their domain structures along with the detection frequency. Notably, for SLCs, the cytoplasmic or extracellular parts were predominantly detected rather than the transmembrane domain. Furthermore, sequence coverage appeared to be highly dependent on the lysis buffer; for instance, SLC12A6 and SLC12A7 exhibited the highest coverage with SDS, while SLC1A4 demonstrated the highest coverage with NP40 (**Fig. 6C**). The same analysis was performed for histones; however, we encountered challenges due to stretches rich in lysine (K) or arginine (R), hindering the generation of suitable tryptic peptides (**Fig. 6D**). Nevertheless, we identified domains with relatively low basic amino acids. In line with our observations for the transmembrane transporters, histones tended to have better sequence coverage with urea and SDS lysis buffer, yet for HIST1H4A urea lysis buffer is of advantage. For a comprehensive picture of all Mix samples, we assessed the sequence coverage of all proteins for each cell type. The average sequence coverage was found to be 20%, with urea yielding the highest sequence coverage in PMH and AS-E2 cells, while NP40 appeared optimal for HepG2 cells MLT (Fig. 6E). Thus, the choice of the best suited lysis buffer not only depends on the context, but also on the specific protein class or sample type of interest in particular if a particular sequence region is of relevance.

### Optimized Lysis Conditions Allow Global Proteomics and Metabolite Affinity Enrichment from the Same Lysate

The investigated lysis conditions were all optimized to generate the highest lysate quality for global proteomics approaches. However, besides direct detection of proteins in lysates with highest coverage by mass spectrometry based global proteomics, it is of interest to study protein complex formation or even interaction of proteins with metabolites by affinity enrichment [29]. For affinity enrichment the three-dimensional, native structure of proteins is of high importance. Therefore, we built on previous knowledge that NP40 lysis buffers containing 0.1 % of SDS have been successfully used in the past for affinity enrichment and employ our optimized NP40 lysis buffer for the lysis of mammalian cell lines AS-E2 and HepG2 [24, 30]. To test for potential intersection points between metabolism and the intracellular protein network, we subjected the lysates of both cell lines to affinity enrichment with inositol hexakisphosphate (InsP_6_). The ligand was immobilized *via* a biotinylated PEG-5-linker to streptavidin coated agarose beads and non-derivatized streptavidin-coated beads were handled in parallel as control experiments [19, 31]. Following protein enrichment, the bound fraction was eluted with excess of soluble InsP_6_ and subjected to SP3-based in-solution digestion. UHPLC-MS/MS analysis, protein search in MaxQuant and data analysis in Perseus revealed the enriched proteins (**Fig. 7A**) [22]. We identified 394 proteins significantly enriched with InsP_6_ in the HepG2 lysate and 314 proteins in the AS-E2 cell lysate (log_2_ fold change = 2, –log_10_(p-value) = 2) compared to the respective control beads (**Fig. 7B**) [32]. An overlap of 193 proteins in both cell lines was observed, which is consistent with the expectation that protein abundance, and therefore also protein enrichment, differs between distinct cell lines (**Fig. 7C**). In line with our expectations, the inositol polyphosphate metabolizing enzymes IP6K, PPIP5K2 and NUDT3 were identified, as well as the known interactors ESYT1 and OCRL, demonstrating the suitability of the NP40 lysis buffer to study metabolite-protein interactions by affinity enrichment (**Fig. 7B**). The identification of known interacting proteins motivated us to analyze the proteins that exhibited the strongest enrichment (the top twenty proteins, with highest fold-enrichment and largest statistical significance, **Fig. 7D**). The majority of these proteins had previously been identified as InsP_6_ interacting proteins in HEK 293T and HCT116 cell lysates. Also, the overlap between the cell lines AS-E2 and HepG2 was high. Here, the most highly enriched proteins were phosphatidyl inositol metabolism-associated proteins (PLCB4, PIP4K2B, PIP4K2A and PREX1)[33–35]. In addition, the serine/threonine kinase CDC42BPA, bearing a PH domain, bound strongly to InsP_6_. Finally, the highly disordered, leucine-rich protein LRRFIP1 appeared to be a relevant target, since its alpha fold structure reveals many basic binding stretches, which are prone to bind inositol polyphosphates. Finally, contrary to previous affinity enrichment procedures, we also identified over 110 plasma membrane-related proteins in the current datasets. Notably, integral membrane proteins, that are particularly hard to lyse, were among these. Nine integral membrane proteins could be found in HepG2, e.g. interferon alpha receptor 1, while seven were detected in AS-E2, including SLC-family members SLC39A6 and SLC12A6. These findings highlighting the broad utility for the NP40 lysis buffer.

**Figure 7:**
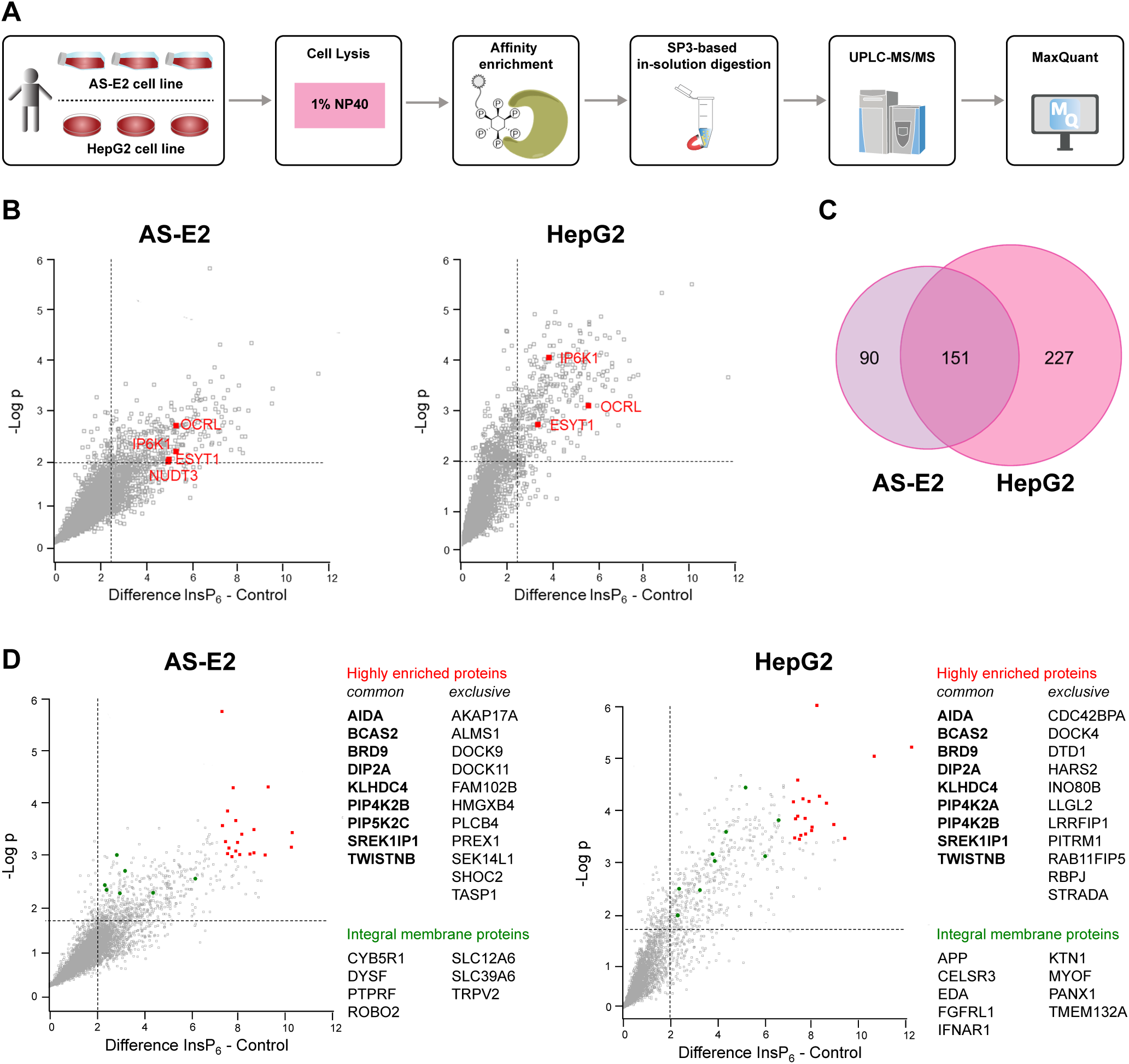
Compatibility of lysis conditions with hexakisphosphate affinity enrichment. **(A)**: Affinity enrichment using HepG2 and AS-E2 cell lysates for identification of binding proteins. The cell lysates were prepared using NP40 lysis buffer and the soluble fractions were used. Biotinylated inositol hexakisphosphate was immobilized on streptavidin-coated agarose beads and applied to both lysates. The full chemical structure can be found in the recent publication by Couto *et al.*, P depicts -OPO ^2-^ rests. **(B)**: Volcano plots showing InsP_6_ binding proteins in AS-E2 and HepG2 lysates after a t-test (log_2_ fold change = 2, –log_10_(p-value) = 2). Known interactors IP6K1, NUDT4, ESYT1 and OCRL were highlighted in red. **(C)**: Venn diagram comparing the significantly enriched proteins by InsP_6_ in AS-E2 and HepG2 lysate. **(D)**: Top 20 proteins that exhibited the strongest enrichment and largest statistical significance in both datasets were highlighted in red and their gene names were depicted beside them. Bold names were found in both datasets within the top 20 proteins. Transmembrane proteins significantly enriched were highlighted in green.

In sum, the results demonstrate that the optimized NP40 lysis buffer is compatible with a previously developed affinity enrichment protocol. This opens possibilities for proteome-wide studies of the interaction between metabolites such as inositol polyphosphates and putative binding proteins, including membrane proteins, which warrant further exploration to e.g. disentangle intersections between signal transduction and metabolism.

## Discussion

Cell and tissue lysis is a critical foundation in proteomics, playing a significant role in shaping the success of LC-MS analyses. However, the efficient extraction of proteins of interest from cells and tissues is not a straightforward process. In proteomics studies, optimizing the lysis process often involves the incorporation of chaotropic agents and detergents into lysis buffers. The roles of chaotropic agents (e.g., urea, guanidine hydrochloride) and detergents (e.g., NP40, SDS) in protein cellular/tissue lysis is to disaggregate large complexes, solubilize membrane-bound species, and denature larger macromolecules to enhance protein extraction[26]. Chaotropic agents disrupt hydrogen bonds and thus enhance protein unfolding during lysis. NP40, for example, is a non-denaturing, mild lysis agent, suitable for isolating cytoplasmic proteins but as it does not disrupt the nuclear membrane is not the detergent of choice for the analysis of nuclear proteins [24]. SDS, in turn, is a strong lysis agent and is effective for the majority of cell types, however it is not well suited for the extraction of sensitive proteins and protein complexes due to its strong denaturing properties [24]. This knowledge guides the incorporation of chaotropic agents or detergents into lysis buffers, aiming for a balance between sufficient denaturation for effective digestion of samples for mass spectrometric analysis and maintaining the integrity of samples for a faithful representation of protein abundances.

Thus, a particular protocol should be unbiased with respect to protein abundance, molecular weight, hydrophobicity, and class. However, to date, the choice of reagents included in lysis buffers and workflows in proteomics is dictated by their orthogonality with downstream LC-MS analysis. In classic in-solution digestion (ISD), chaotropic agents or detergents have to be removed or diluted before (trypsin activity) or latest after protein digestion to ensure compatibility with LC-MS analysis, in order to prevent interference with separation and detection. Cleanup methods, such as Stage-tip and filter-aided sample preparation (FASP), are widely adopted, but these protocols are complex and time-demanding, frequently leading to poorly quantifiable data. Furthermore, specific reagent compatibilities pose challenges; some cleanup approaches may not align with reagents like NP40 and SDS, which are detergents of choice in signal transduction studies as they enhance cell lysis efficiency, as they can be problematic due to their interference with chromatography and electrospray ionization during LC-MS analysis. Recent advancements, such as single-pot, solid-phase-enhanced sample-preparation (SP3) techniques, have overcome these challenges [11, 36]. SP3 allows the use of previously deemed unsuitable reagents for mass spectrometry-based proteomics, representing a paradigm shift in sample preparation methodologies. Our study systematically evaluated different reagents widely incorporated into lysis buffers using SP3 for sample preparation, focusing on proteomes from different biological samples, including mouse tissue and various cell lines as well as primary cells. The lysis buffers tested included NP40, SDS, GdnHCl, and urea, each at their commonly used concentrations. These reagents were added to a variation of RIPA lysis buffer, which is the most commonly used buffer for total protein extraction from mammalian cells and tissues. Classical RIPA lysis buffer comprises a low concentration of SDS, deoxycholate for disruption of protein-protein interactions, and other components [37]. In our study, we replaced the conventional low concentration of SDS with individually tested reagents. Moreover, we implemented a dual-sample lysis approach, analyzing Supernatant and Pellet fractions separately and in combination (Mix). This strategy provided a comprehensive understanding of how each reagent influenced various aspects of proteomic data quality and quantification. Following a successful lysis protocol and protein digestion, samples were analyzed in an LC-MS system. Observing the chromatograms, we tracked the peptide elution profiles and observed distinct differences emerging among various reagents. NP40, SDS, and urea exhibited a comparable profile. As anticipated, the utilization of the SP3 method resulted in the absence of NP40 or SDS traces in the peptide solutions, as evidenced by the chromatogram consistency of the samples of the NP40 or SDS lysis buffers with the urea-containing lysis buffer. Such consistency also indicates that the tryptic digestion was not affected by one or the other reagents as SDS and urea can interfere with enzymatic digestion, resulting in reduced proteolytic efficiency [26].

However, the distinctive chromatogram profile of samples from GdnHCl lysis buffer, especially for AS-E2 cells and PMH, raises questions about the efficiency of these lysis buffers in specific cellular contexts. Additionally, the evaluation of proteome coverage provided insights into the reliability of each lysis buffer in the extraction and identification of proteins. While all lysis buffers demonstrated similar performance concerning the number of identified proteins in supernatants, the suboptimal performance of GdnHCl, particularly in AS-E2 cells, underscores the necessity for careful consideration of lysis buffer selection. On the contrary, Neset et al. [6] reported that utilizing either an SDS- or GdnHCl-based lysis buffer performed comparably when either lysis buffer was combined with SP3 for LC-MS analysis of HeLa cells. Furthermore, GdnHCl has been acknowledged as one of the most efficient chaotropic agents for protein extraction from extracellular matrix (ECM)-rich tissues, such as tendon and bone [8, 38, 39]. This observation emphasizes the critical importance of understanding the intricate interplay between the lysis conditions and distinct cell/tissue types before designing an LC-MS-based proteomics analysis. The consistent efficacy of individual supernatants in proteome extraction across diverse sample types resonates with studies highlighting the importance of efficient lysis in obtaining comprehensive protein profiles [40]. In addition to that, an efficient lysis yields proteins from all the cellular compartments in an unbiased manner.

Extracting proteins from the plasma membrane and nucleus is more difficult than the extraction of cytosolic proteins. One major problem encountered in solubilizing membrane proteins is related to the hydrophobicity of the transmembrane domain of the proteins. Unlike membrane-associated proteins that can be removed via changes in ionic strength or pH, extraction of integral membrane proteins, especially with multiple transmembrane domains, normally requires harsh solubilizing conditions as they tend to aggregate and precipitate in aqueous solution [41, 42]. Nuclear proteins, in turn, are moderately soluble [43], but they are tightly packed in the nuclear structure, strongly interacting with the chromatin structure. Breaking down these structural barriers to release proteins for analysis requires effective lysis reagents. To investigate potential cellular component biases intrinsic for each lysis buffer, we focused on the individual analysis of supernatant and pellet samples of the different lysis buffers employed. Interestingly, our results indicate a considerable presence of proteins, both in terms of quantity and intensity, within the pellet fractions across various tested lysis buffers. In this case, the conventional practice of discarding the pellet, would lead to a significant loss of numerous proteins, critically compromising the interpretation of the results, both qualitatively and, most critically, quantitatively. Furthermore, our studies revealed disparities in protein composition and distribution between the supernatant and pellet fraction. Particular examples are the solute-carrier (SLC) superfamily of membrane transport proteins that constitute a diverse family of membrane proteins engaged in transporting nutrients, metabolites, xenobiotics, and drugs. Although they are present throughout the cell, residing in the membrane of nearly every organelle, a majority are situated in the cell membrane. Their transmembrane structure is characterized by hydrophobic alpha-helices interconnected by hydrophilic intra- and extra-cellular loops [44–46]. Likewise, histones were selected to serve as representatives of nuclear proteins. Histones are categorized into five families: H1/H5 (linker histones), H2, H3, and H4 (core histones). The nucleosome core is composed of two H2A-H2B dimers and a H3-H4 tetramer. The extensive wrapping of DNA around histones primarily results from the electrostatic attraction between the positively charged histones and the negatively charged phosphate backbone of DNA [27]. This close association may pose challenges in accurately representing histones in proteomics analyses. For many year, NP40 lysis buffer have been employed for the preparation of cellular extracts for antibody based assays such as immunoblotting, ELISA, or multiplexed bead based arrays as well as the analysis of protein complexes e.g. by immunoprecipitation, due to its properties as a milder detergent capable of solubilizing proteins without strong denaturation. Taking HepG2 as an example, our results reveal a distinct cellular component pattern associated with the NP40 lysis buffer: a prominent presence of plasma membrane proteins in the supernatant fraction and an increased abundance of nuclear proteins in the pellet fraction for all sample types. Moreover, a differential expression analysis comparing the supernatant and pellet of the NP40 lysis buffer showed that 1209 proteins were most abundant in the supernatant fraction, with SLCs emerging as some of the most abundant proteins. Conversely, 180 proteins, including histones, were observed to be more abundant in the pellet fractions. Overall, our data revealed significant distinctions between the protein composition of supernatant and pellet fractions across all lysis buffers and sample types, challenging conventional practices and advocating for a dual-sample lysis procedure when using NP40 lysis buffers.

However, it is important to note that adopting this method and routinely analyzing supernatant and pellet fractions separately would result in doubling the LC-MS analysis time. We demonstrate that a potential strategy to overcome this challenge involves combining the supernatant and pellet fractions at the sample level immediately after the double-lysis procedure and before the LC-MS analysis. Moreover, our data reveals, in the case of the use of NP40 lysis buffer, that the mix approach not only closely mirrors the composition of individual supernatant and pellet fractions in terms of the total number of identified proteins but also effectively restores abundances, as e.g. observed in the cases of SLCs and histones. Among all the tested reagents, SDS stood out as exhibiting a balanced and uniform distribution of proteins in both the supernatant and pellet fractions. This underscores its versatility in lysing proteins from diverse cellular compartments, a characteristic previously discussed and documented [47]. A comparable trend was observed for urea lysis buffer; however, though not as pronounced as NP40, urea demonstrated a slight inclination to overrepresent a specific class of proteins (nuclear) in the supernatant fraction. These findings align with the ongoing discourse regarding the existence of a truly universal protocol with minimal or no extraction bias, applicable across all sample types [48–50].

From a reagent perspective, we have highlighted biases, advantages, and pitfalls associated with a selected range of reagents. This emphasizes the necessity for a reassessment of conventional practices to enhance the adaptability of lysis buffers to the specific requirements of individual proteomics studies. Furthermore, our findings indicate that the selection of a lysis buffer significantly impacts the sequence coverage of specific proteins. In the context of the plasma membrane category, our analysis identified proteins exhibiting statistically significant differences in sequence coverage. Notably, over 400 plasma membrane proteins displayed varying sequence coverage, with a particular emphasis on SLCs. SLCs, our results highlighted a prevalence of cytoplasmic or extracellular domains over the transmembrane region. Intriguingly, certain SLCs exhibited optimal coverage with SDS, while others achieved the highest coverage with NP40. Similarly, within the nuclear category, our investigation revealed more than 1000 proteins manifesting divergent sequence coverage, with histones being prominently featured. Moreover, the investigation of histones was more challenging due to the number of lysine (K) or arginine (R)-rich stretches hindering the generation of suitable tryptic peptides. However, in concordance with our observations, histones demonstrated enhanced sequence coverage when utilizing the urea lysis buffer. The comprehensive overview of the sequence coverage of all Mix samples of all cell types, showed that on average, the sequence coverage of proteins was most comprehensive with urea for PMH and AS-E2, while NP40 lysis buffer demonstrated optimal results for MLT. This implies that the selection of a lysis buffer should be tailored to the specific protein class or sample type, particularly when targeting specific sequence regions of interest. These findings underscore the critical link between lysis buffer choice and sequence coverage, influencing the suitability of peptides for proteomics. Since peptides play a central role as protein surrogates, our observations highlight the need for an informed and adaptive approach in the selection of proteotypic peptides [51]. This becomes crucial for achieving unequivocal identification and quantification of (targeted) proteins, emphasized in quantitative and targeted proteomics analysis.

When planning an experiment, one has to keep in mind what purposes a sample shall serve. Often, samples are rare and only little material is available, e.g. in case of patient tissue, and it is of advantage if the same sample can be used for various analyses. In the case of cellular or tissue lysis samples might not only be prepared for LC-MS, but the lysates could also first be utilized to explore protein-protein interactions or protein-metabolite binding through immunoprecipitation or affinity enrichment assays. In these cases, the native state of proteins has to be maintained and thus a lysis buffer has to be chosen not only securing high quality LC-MS measurements, but also leaving the three-dimensional structure of proteins intact. Since SDS, urea and GdnHCl interfere with the 3D structure of a protein, of the four lysis buffers investigated only the NP40 lysis buffer is compatible with techniques requiring the native state of a protein [52]. We demonstrated in an affinity enrichment with InsP_6_ that indeed NP40 lysis buffer is suitable for detecting protein interaction partners. The large number of proteins of enriched proteins, especially known InsP_6_ interacting proteins, such as IP6K1 or NUDT3 [53], prove that the NP40 lysis buffer is not only suitable for reliable LC-MS measurements, but also for native protein interaction techniques like affinity enrichment with small molecules such as InsP_6_. Our results demonstrate that by employing the NP40 lysis buffer and affinity enrichment followed by mass spectrometry based interactome characterization not only confirms previously established intracellular interactions but extents the spectrum to integral membrane proteins thus opening an entirely novel layer to study interactions points in cellular networks e.g. between metabolism and signal transduction.

These findings show exemplarily that the choice for lysis conditions depends on many factors and that there is not a single best solution. Yet with the knowledge we provide a well-founded choice can be made.

## Conclusion

In conclusion, our comprehensive investigation into the influence of lysis buffers on the outcome of proteomics studies has shed light on crucial aspects for mass spectrometry analysis. The careful examination of diverse biological samples using a variety of lysis buffers provide insights into their differential impact on protein extraction, denaturation, and subsequent peptide analysis. Our findings underscore the significance of selecting appropriate lysis buffers tailored to specific sample types. Notably, SDS emerge as a consistently reliable lysis buffer, exhibiting stable and reproducible performance across diverse cellular contexts. Additionally, our results encourage a dual-sample lysis approach, challenging the conventional practice of analyzing only the Supernatants post-lysis. Moreover, our results suggest that the efficiency gained from consolidating Supernatant and Pellet fractions in a Mix sample and combining them into one LC-MS run, as opposed to conducting two separate runs, does not compromise the quality of analytical insights. Furthermore, the evaluation of protein sequence coverage emphasizes the importance of considering the intricacies of each lysis buffer for optimal results, especially if specific protein domains must be represented. The same is crucial when selecting peptides for targeted proteomics. The variation in sequence coverage across different lysis buffers and protein categories (plasma membrane and nucleus) highlights the need for a thoughtful process selection based on the characteristics of the specific sample under investigation. In practical terms, our research provides valuable guidance for refining bottom-up proteomics workflows. By tailoring lysis buffer selection to the unique attributes of biological samples, researchers can enhance the robustness, reproducibility, and efficiency of their mass spectrometry analyses. Furthermore, we demonstrate that a mild lysis with NP40 lysis buffer not only allows efficient mass spectrometry-based analysis of integral membrane proteins but also interactions studies of proteins with metabolites. Thus, this study contributes not only to the optimization of protein extraction protocols but also challenges traditional approaches, offering a more streamlined and effective strategy for future proteomic investigations and studies of interactions between metabolism and proteins.

## Acknowledgements

We thank Franziska Gödtel, Yannik Dieter, Alexander Held, Sandra Bonefas and Xiaomeng Li for their technical assistance.

## Data Availability

The original MS raw data files and the MaxQuant search result files are available on the ProteomeXchange consortium via PRIDE (https://www.ebi.ac.uk/pride/).

